# Gut microbiota features of the geographically diverse Indian population

**DOI:** 10.1101/478586

**Authors:** Sudarshan A. Shetty

## Abstract

Population-level microbial profiling allows for identifying the overarching features of the microbiome. Knowledge of population specific base-line gut microbiome features is important due to the widely reported impact of geography, lifestyle and dietary patterns on the microbiome composition, structure and function. Here, the gut microbiota of more than 1000 subjects across the length and breadth of India is presented. The publicly available 16S rRNA gene profiling data of faecal microbiota from the Landscape Of Gut Microbiome - Pan-India Exploration (LogMPIE) study representing 14 major cities, covering populations from northern, southern, eastern and western part of India analyzed. Majority of the dominant OTUs belonged to the Firmicutes, Bacteroidetes and Proteobacteria phyla. The rarer fraction was comprised of OTUs mainly from the phyla Verrucomicrobia and Spirochaetes. The median core size was estimated to consist of 12 OTUs (>80% prevalence) dominated by representing genera *Prevotella*, *Faecalibacterium, Bacteroides, Roseburia, Megasphaera*, *Eubacterium* and *Gemmiger*. Geographic location explained majority of the variation in the gut microbiota community structure. The observations of the present study support the previous reports of *Prevotella* dominance in the Indian population. The *Prevotella*/*Bacteroides* ratio was high for the overall population irrespective of geographic location and did not correlate with BMI or age of the participants. Despite a rapid transition towards a western lifestyle, high prevalence of *Treponema* in the Indian gut microbiota suggests that the urban population still harbors signatures of the traditional gut microbiome. The results presented here improve the knowledge of baseline microbiota in the Indian population across the length and breadth of the country. This study provides a base for future studies which need to incorporate numerous other confounding factors and their impact on the observed characteristics of the Indian gut microbiome.

## Introduction

Numerous population-level studies have been conducted to investigate base-line as well as population specific characteristics of the human gut microbiome. These included human populations from the USA, Netherlands, Belgium, Denmark, Spain, Africa, Venezuela, China, Mongolia, Fiji, Israel and Papua New Guinea (Qin *et al.*, 2010, Jalanka-Tuovinen *et al.*, 2011, Huttenhower *et al.*, 2012, Qin *et al.*, 2012, Yatsunenko *et al.*, 2012, Lahti *et al.*, 2014, Li *et al.*, 2014, Zhang *et al.*, 2014, Martínez *et al.*, 2015, O’Keefe *et al.*, 2015, Yano *et al.*, 2015, Falony *et al.*, 2016, Rothschild *et al.*, 2018). These studies have uncovered a vast diversity of the gut microbial communities as well as identified several factors influencing the microbiome, including age, ethnicity, dietary patterns, geographical location, consistency of faecal samples (Bristol stool chart), lifestyle, etc. It is commonly observed that *Bacteroides* is associated with high protein diet while *Prevotella* is associated with high fibre diet (David *et al.*, 2014, Gorvitovskaia *et al.*, 2016). Several bacteria have been identified as part of the core microbiota in diverse populations as well as common core functions have been reported (Turnbaugh *et al.*, 2009, Jalanka-Tuovinen *et al.*, 2011, Huse *et al.*, 2012, Li *et al.*, 2014, Falony *et al.*, 2016). These studies have directed mechanistic studies and clinical trials for identifying health and disease related diagnostic biomarkers and development of strategies for modulation of the microbiome for health benefits (De Filippo *et al.*, 2010, Cotillard *et al.*, 2013, David *et al.*, 2014, Schubert *et al.*, 2014, Zeller *et al.*, 2014, O’Keefe *et al.*, 2015, Baxter *et al.*, 2016, Desai *et al.*, 2016).

However, similar information on population-level characteristics of the gut microbiota in a Indian subjects with representative sampling across its geography are limited (Ghosh *et al.*, 2013, Shetty *et al.*, 2013, Dehingia *et al.*, 2015, Bhute *et al.*, 2016). Previously, the importance of understanding the complexity and diversity of the gut microbiome in the Indian population was reviewed (Shetty *et al.*, 2013). Several features that make the subjects in the Indian sub-continent different such as dietary habits, socio-economic situations, societal traditions of dietary habits, vast genetic diversity as well as prevalence of diseases not associated with altered gut microbiome was documented (Shetty *et al.*, 2013). The YY-paradox is an important differentiating factor of human populations in the Indian sub-continent, where Indians with same body mass index as a Western individual have three times the fat content (Yajnik & Yudkin, 2004). This makes the application of BMI to classify obese and non-obese status debatable for the Indian population (Yajnik & Yudkin, 2004, Shetty *et al.*, 2013). A first step towards better understanding the role of gut microbiome on health is to catalogue the population specific microbial diversity, composition and structure using a large representative sample. This “stamp-collection” process has been a driving factor for several of the currently known disease and health associations and development of potential microbiome biotherapeutic candidates (Qin *et al.*, 2012, Everard *et al.*, 2013, Lahti *et al.*, 2014, Dao *et al.*, 2015, Falony *et al.*, 2016, Plovier *et al.*, 2017, Shetty *et al.*, 2017). These features can be further linked to several populations specific features as well as individual-specific microbiota features using extensive phenotyping and measurement of environmental covariates (Falony *et al.*, 2016).

Here, results from the largest standardized collection of the gut microbiota profiles of heterogeneous Indians subjects across geography is presented. The primary focus of the study was to identifying compositional variation, similarities and dissimilarities in gut microbial community structure and identifying the core microbiota across geographic landscape. Furthermore, the underlying variation across the gut microbial community structure was found to be associated with the *Prevotella*/*Bacteroides* ratio.

## Results and Discussion

### Brief description of the study population

The detailed the subject data is described in the original article reporting the LogMPIE study (Dubey *et al.*, 2018). Briefly, the study reported microbial profile of 1004 Indian individuals residing in 14 cities different cities. These broadly covered the populations representative of north, west, east and south geographic areas of the country. Data on lifestyle, body mass index (BMI), age and gender were reported. The mean and standard deviation for age was 37.2 ± 11.9, for BMI was 27.9 ± 4.9 represented by 420 females and 584 males. Out of the 1004 subjects, 556 were categorised as non-obese and 448 as obese based on the BMI. The subjects were further categorised following a sedentary and non-sedentary lifestyle. The metadata was limited to these factors and other important metadata such as dietary intake (vegetarian/non-vegetarian, ratio of carbohydrates to protein in diet, consumption of yogurt with live bacterial cultures, etc.), stool consistency, history of medications was not reported. Therefore, the preliminary analysis here does not address the effects and/or contribution of these factors to the variation in gut microbiota.

### Microbial composition and community level variation across geography

The microbiota composition showed differences at phylum level in individuals from the different geographic zones (Figure 1). Prominent differences in relative abundance were observed in the phyla Bacteroidetes, Firmicutes and Proteobacteria. Individuals from east and north harboured higher abundances of Bacteroidetes compared to west and south. Individuals from north and south harboured relative higher abundance of Spirochaetes (Figure 1).

**Figure 1:**
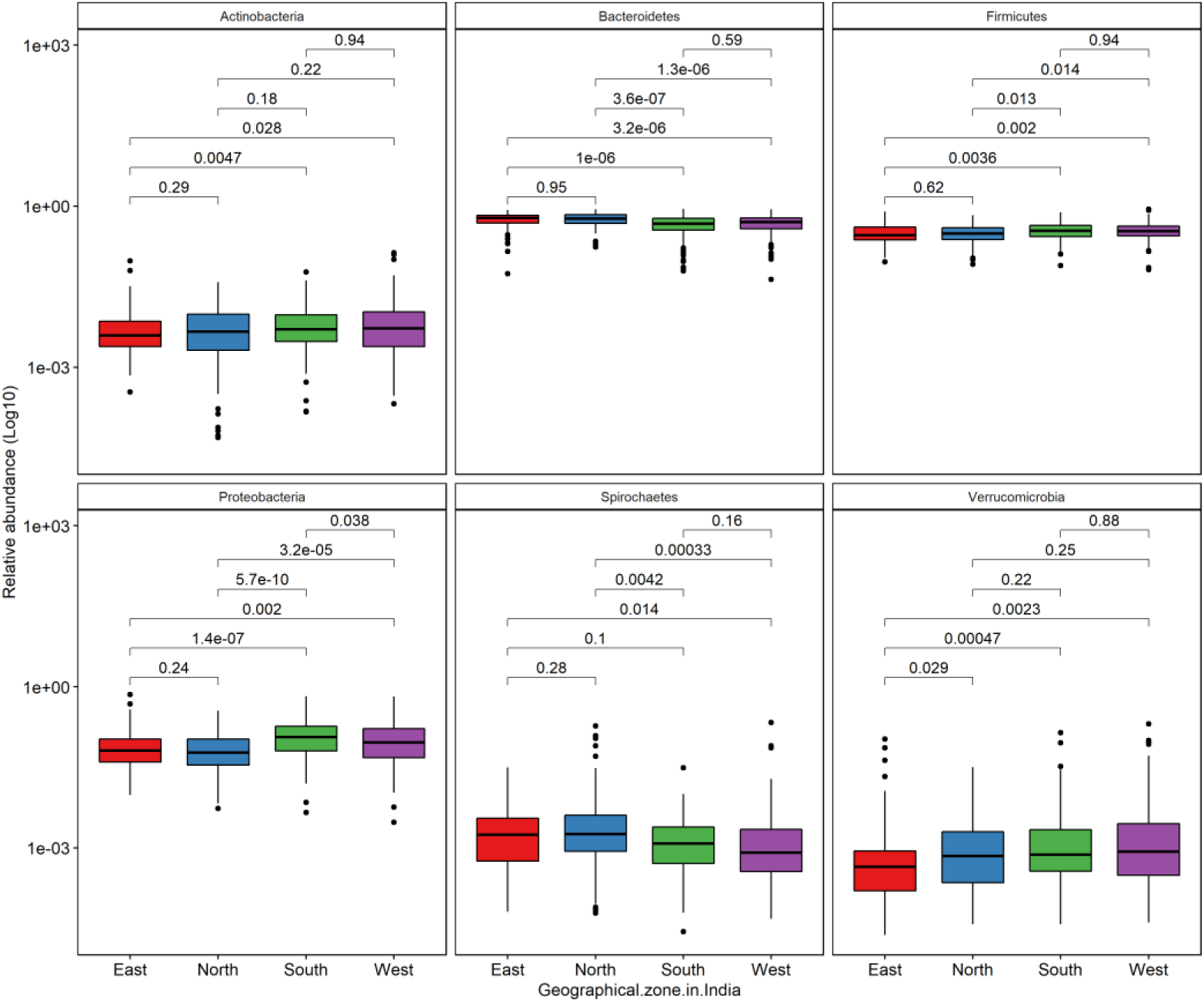
Comparison of relative abundances of major phyla in the gut microbiota of Indians. The p-values were calculated using Wilcoxon test.

At genus level, *Prevotella* was abundant across the geographic landscape, followed by *Faecalibacterium* (Figure 2). Genus *Bacteroides*, *Megasphaera*, *Parasutterella*, *Haemophilus* showed variable abundances, where few individuals had more than 0.4 (proportional) abundance. Comparison of microbiota of Indians with other populations has reported the enrichment of *Prevotella* and *Megasphaera* (Bhute *et al.*, 2016). The observation in a large population here provides further support for their association with Indian gut microbiota. *Megasphaera* is a butyrate and propionate producer both of which are known for anti-inflammatory properties (Hosseini *et al.*, 2011, Lin *et al.*, 2012, Louis & Flint, 2017). The observation of variable abundances of *Parasutterella* and *Haemophilus* is intriguing as these are hardly reported to be highly prevalent and/or abundant in gut microbiota of healthy western adults (Human Microbiome Project, 2012, Falony *et al.*, 2016). However, abundance of *Parasutterella* was associated with urban Mongolian microbiota (Zhang *et al.*, 2014). The physiological and metabolic characterization is currently focused on the anaerobic lifestyle of bacteria from Bacteroidetes phyla, Lachnospiraceae and Ruminococcacaea families (Barcenilla *et al.*, 2000, Sonnenburg *et al.*, 2010, Flint *et al.*, 2012, Flint *et al.*, 2012, Reichardt *et al.*, 2014). All of which have been reported to be dominant in the Western population. However, microbiota analysis of non-western populations advocates the need to focus on obligate anaerobic bacteria from phyla Proteobacteria and Spirochaetes to understand their role in health of non-western adults (Martínez *et al.*, 2015, Bhute *et al.*, 2016, Das *et al.*, 2018).

**Figure 2:**
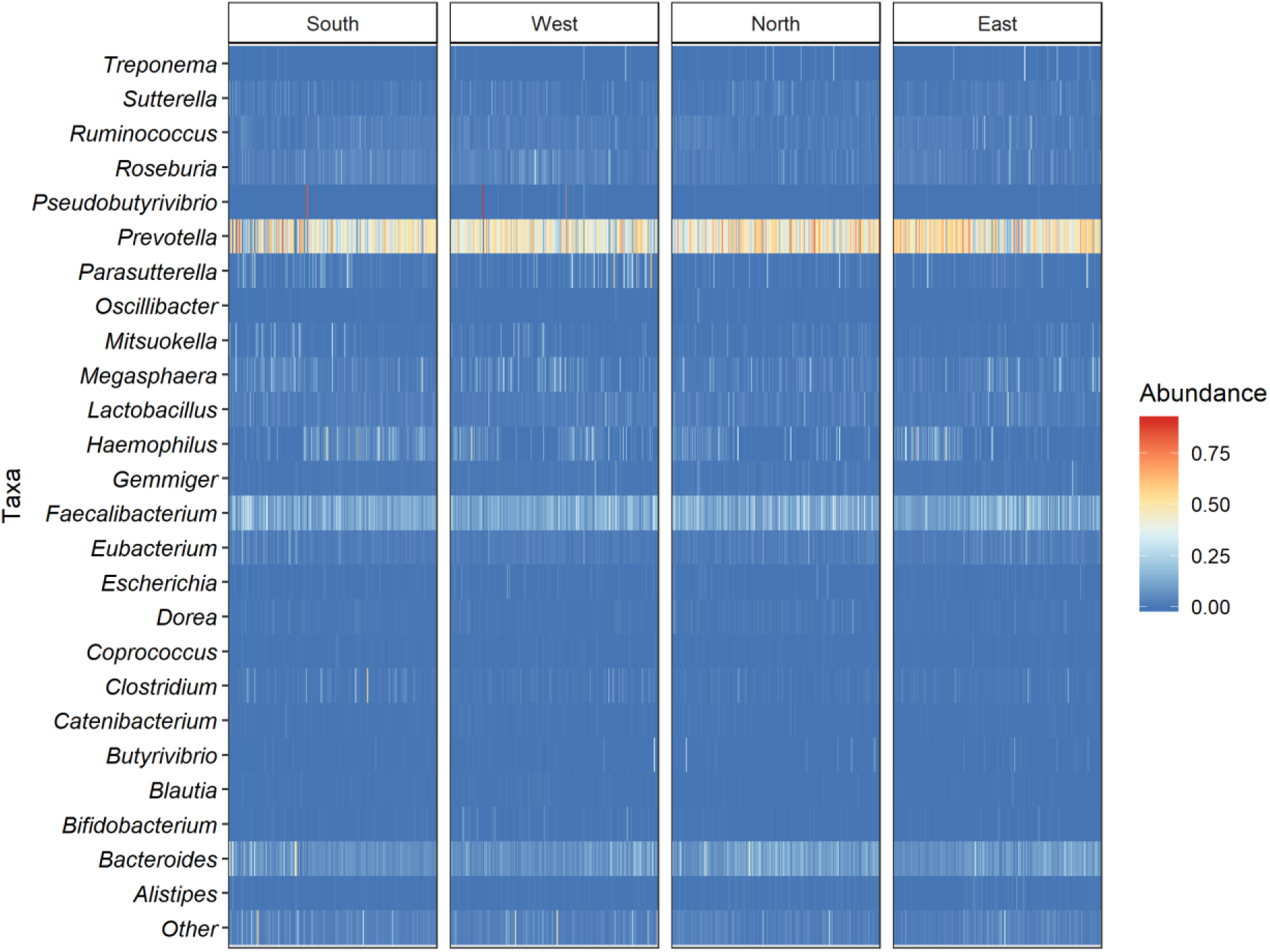
Inter-individual variation in relative abundance of top 25 gut microbial genera in subject from different geographical zones in India.

Based on unconstrained principal coordinate analysis (PCoA) analysis of OTU-level, no major separation was observed between the populations from different broadly classified geographic locations *i.e.* north, west, east or southern part of the country (Figure 3). The microbial community structure was not significantly associated with obesity status of the individuals within the population (PERMANOVA, *P* = 0.585). Geographical location (city of residence) explained the most variation (PERMANOVA, R2=0.10, Pr(>F) = 0.001), followed by geographical zone (PERMANOVA, R2 = 0.02, Pr(>F) = 0.001), gender (PERMANOVA, R2=0.002, Pr(>F) = 0.009) and lifestyle pattern (PERMANOVA, R2 = 0.005, Pr(>F) = 0.001).

**Figure 3:**
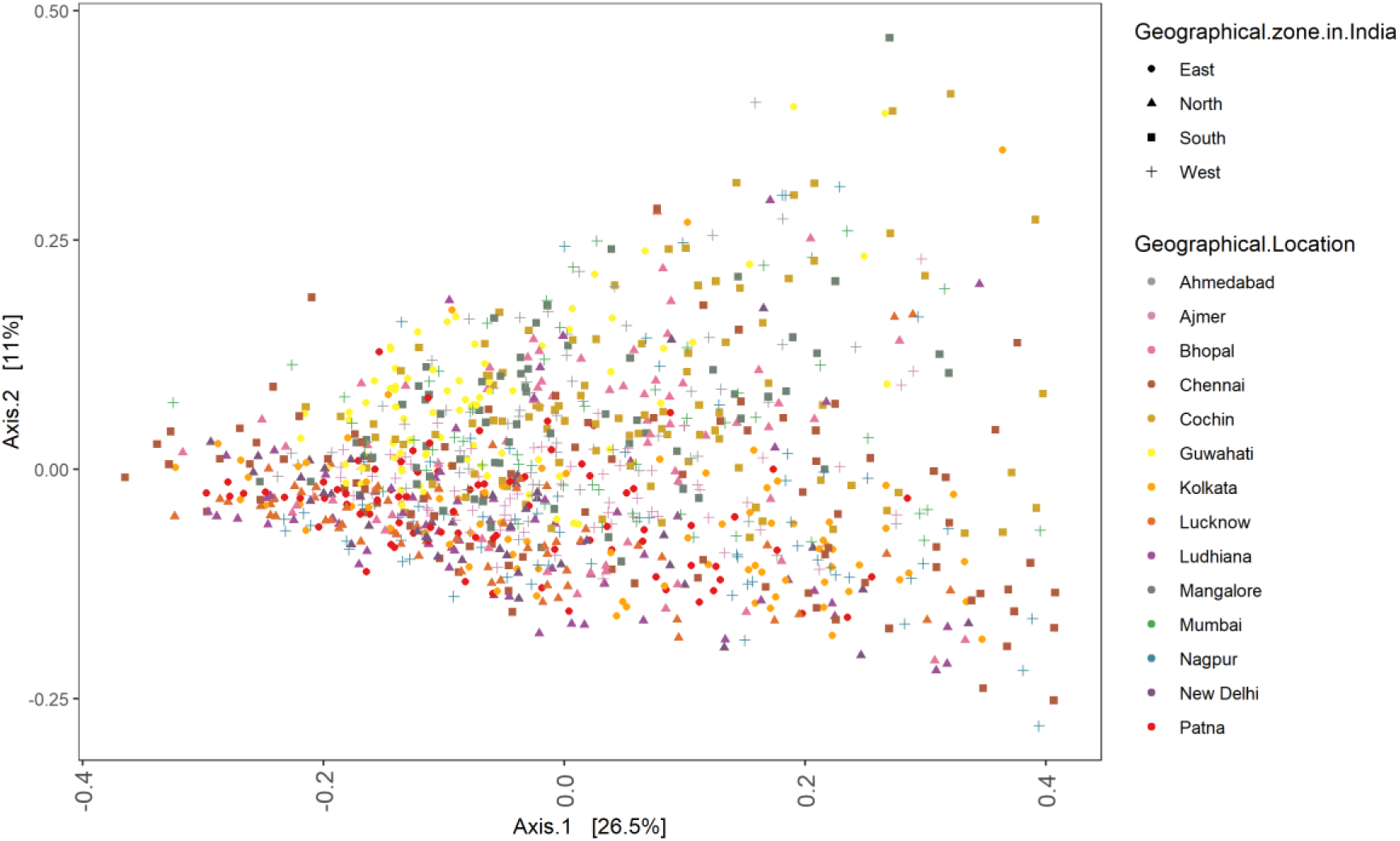
Principal coordinates analysis (PCoA) based on Bray-Curtis dissimilarity based on OTU relative abundances.

Within each of the geographic zones *i.e.* north, south, east and west, there are differences in the microbiota structure between the cities (Supplementary Figure 1). The above observations demonstrate that environmental factors are a major driver of the gut microbiota, especially the location of residence in the cohort investigated in this study. Further highlighting the effect of geographic locations and related confounding factors as an important challenge in identifying health and disease associated biomarkers for the Indian population. A major metadata lacking in the LogMP study is the dietary intake. Each of the cities sampled in the Log MP study is separated at least 200km, while most are separated by a distance of more than 500 km. Each of these cities has distinct lifestyle as well as culinary traditions. A comprehensive characterisation of the gut microbiota and its association with disease will require incorporating information on diet and lifestyle related cofounding factors in future studies.

### Prevalent dominant and rare bacteria in the Indian gut microbiota

Both the dominant and rare fractions of the microbiome play an important role in stability and resilience of the microbial community (Shade *et al.*, 2014, Lynch & Neufeld, 2015, Shetty *et al.*, 2017, Delgado-Baquerizo *et al.*, 2018, Jia *et al.*, 2018). Identifying bacteria that comprise the dominant and rare fractions is important to better understand their potential role and consequent impact on the functioning of the microbiome. Based on the abundance-occupancy analysis, OTUs from phyla Firmicutes, Bacteroidetes and Proteobacteria were identified as covering the abundant fractions in the Indian gut microbiota (Figure 4). The most abundant OTU was from the Firmicutes phyla was *Faecalibacterium prausnitzii* (OTU000444; 0.14, 100%), from Bacteroidetes was *Prevotella copri* (OTU000745; 0.4, 99%), from phylum Actinobacteria was *Bifidobacterium bifidum* (OTU000175; 0.002, 68%), from Proteobacteria was *Haemophilus parainfluenzae* (OTU000484; 0.03, 87%), from Spirochaetes was *Treponema succinifaciens* (OTU000961; 0.004, 73%) and from Verrucomicrobia was *Akkermansia muciniphila* (OTU000067; 0.002, 61%). Three OTUs from Proteobacteria were present in more than 90% of the samples (OTU000703:Parasutterella, OTU000468:Gemmiger and OTU000935:Sutterella with 0.02, 0.01 and 0.009 mean proportional abundance). A detailed list is given in supplementary table 1.

**Figure 4:**
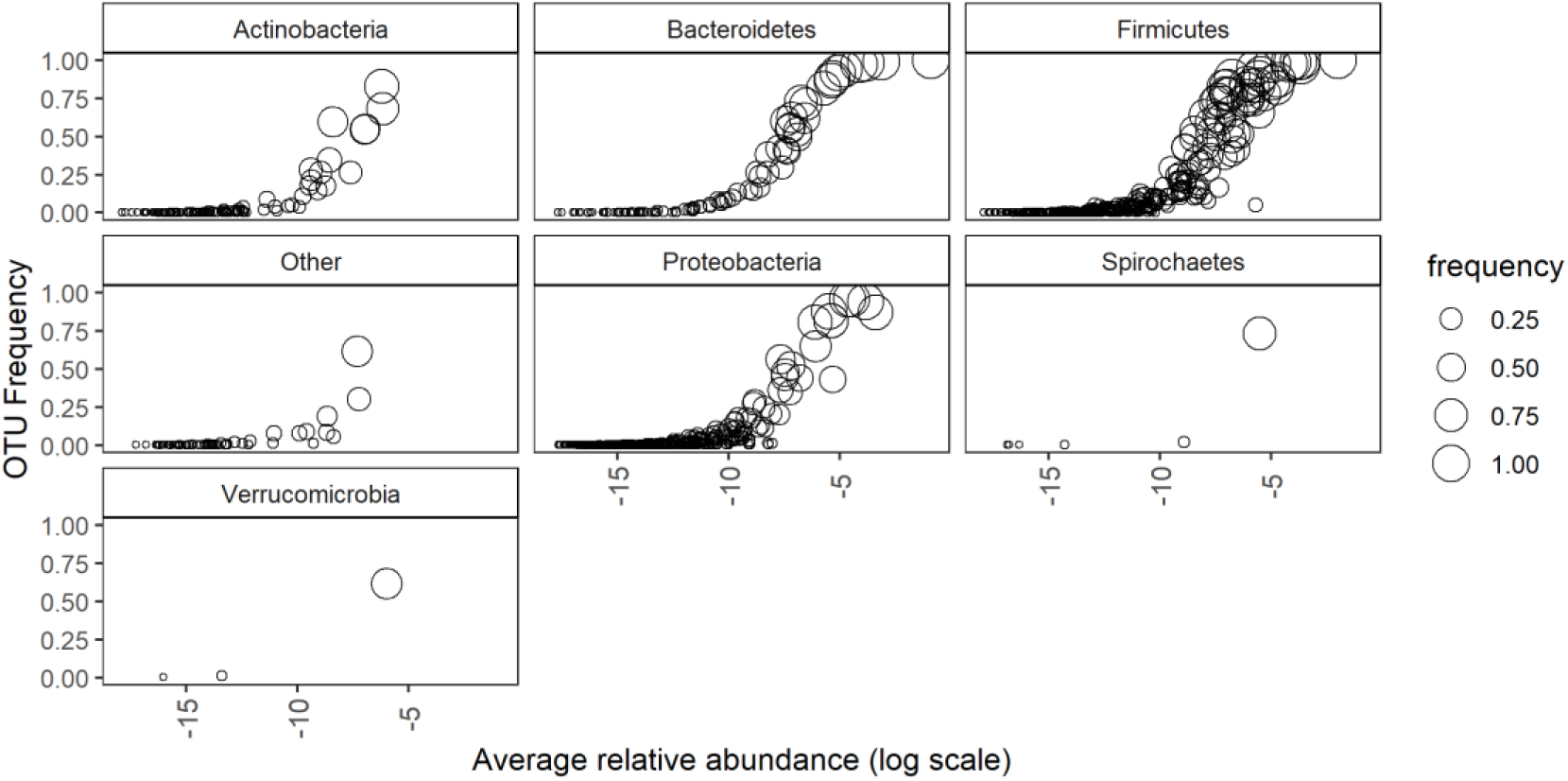
Occupancy-Abundance relationship for OTUs from major phyla in the Indian gut microbiota (n=1003). The x-axis is log transformed for clarity.

In the present study, re-analysis of the data was done to gain detailed insight into the core microbiota following the bootstrap approach as reported previously (Jalanka-Tuovinen *et al.*, 2011, Salonen *et al.*, 2012, Shetty *et al.*, 2017). The change in core size with respect to various abundance and prevalence thresholds is shown in Figure 5A. The median core size and the core OTUs were estimated to consist of 12 OTUs (minimum relative abundance threshold of 0.0001 and presence in at least 80%). These included otu000745 (*Prevotella copri*), OTU000444 (*Faecalibacterium prausnitzii*), OTU000162 (*Bacteroides plebeius*), OTU000756 (*Prevotella stercorea*), OTU000542 (*Lactobacillus rogosae*), OTU000814 (*Roseburia faecis*), OTU000149 (*Bacteroides coprophilus*), OTU000834 (*Ruminococcus gnavus*), OTU000594 (*Megasphaera elsdenii*), OTU000426 (*Eubacterium eligens*), OTU000468 (*Gemmiger formicilis*), OTU000148 (*Bacteroides coprocola*). Investigation of varying abundance and prevalence thresholds for inclusion of core microbiota aided in identifying both abundant and rare members of the core microbiota in the Indian population (Figure 5B). The *Prevotella copri* was identified as the most prevalent and dominant core bacteria across a range of abundance and prevalence thresholds (Figure 5 A and B). This is in accordance with recent report on the gut microbiota of tribal as well as urban Indian populations (Dehingia *et al.*, 2015, Bhute *et al.*, 2016, Das *et al.*, 2018, Tandon *et al.*, 2018). At the genus level, the core microbiota contributed to a large fraction of the total microbiota across geographies and gender (Supplementary figure 2).

**Figure 5:**
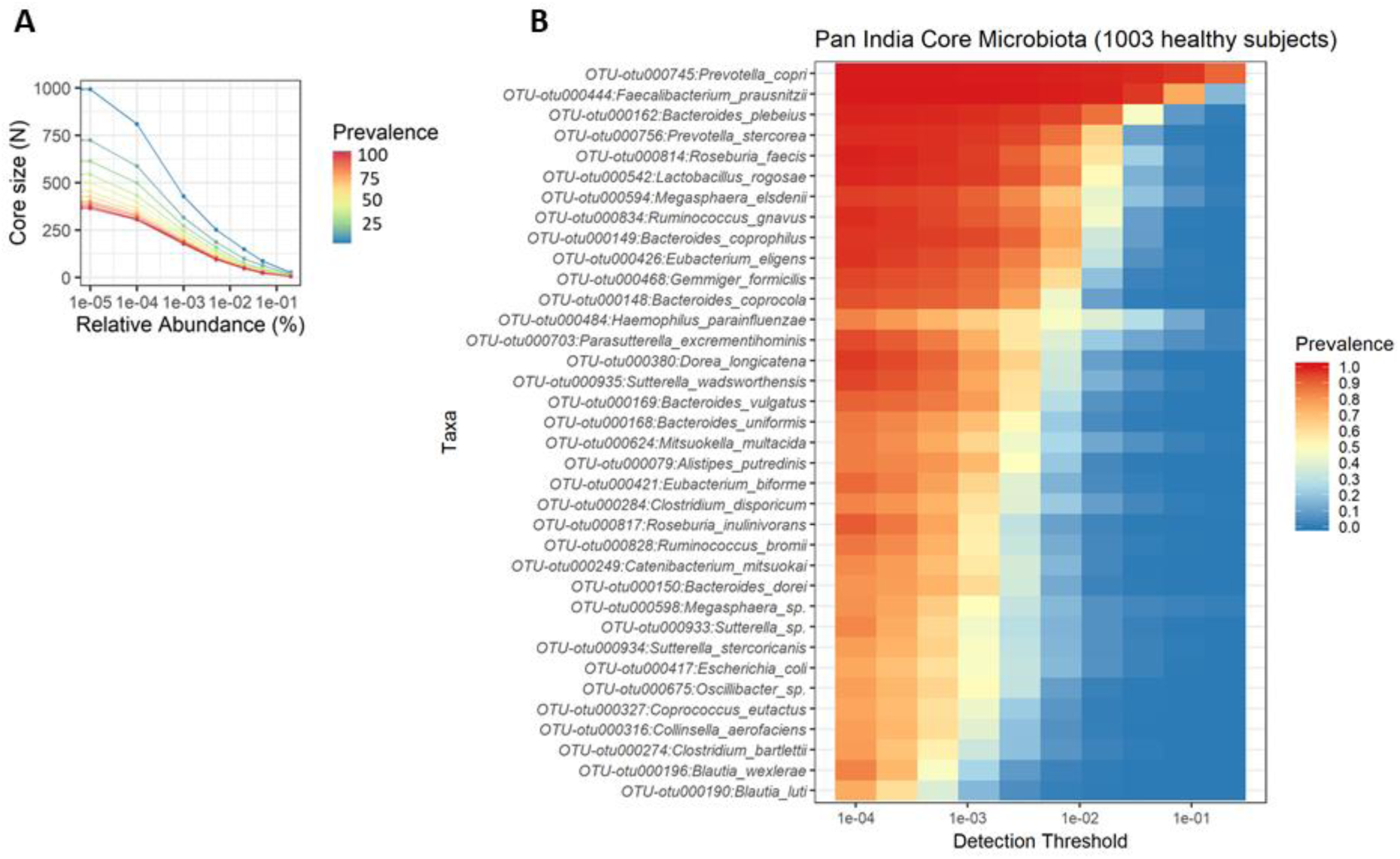
Core microbiota in Indian population. A] The difference in number of core OTUs and their prevalence at different abundance thresholds. B] Heatmap depicting the core OTUs, their prevalence at different detection thresholds (relative abundance).

The dominance and prevalence of *Faecalibacterium* is associated with both western and non-western populations (Falony *et al.*, 2016, Shetty *et al.*, 2017). Prevalence and abundance of *Prevotella* is associated with gut microbiota of non-western populations (Falony *et al.*, 2016). In our study, we identify both of these genera as a part of the Indian core microbiota. These bacteria have a range of metabolic traits related to degradation of complex polysaccharides (David *et al.*, 2014, Heinken *et al.*, 2014). However, there is a lack of direct evidence of complex fibre degradation ability for *Prevotella copri*, the most abundant and prevalent species detected in the gut microbiome. This species is known to have β-Galactosidase, α-Arabinofuranosidase and β-Glucosidase activity (Hayashi *et al.*, 2007). On the contrary numerous evidence exists for polysaccharide degradation ability in species from the genus *Bacteroides* (Sonnenburg et al., 2010). Further investigation of physiology and polysaccharide degrading ability of *Prevotella* and its species/strains across human populations will be crucial to better understand its role in the gut microbiome.

### Prevotella dominance is hallmark of Indian gut microbiota irrespective of geographic location, age, gender and BMI

The dominance of *Prevotella* or *Bacteroides* is an important property of the human gut microbiome as these bacteria are known to be biomarkers of diet and lifestyle (Gorvitovskaia *et al.*, 2016). Hence, the *Prevotella* versus *Bacteroides* (P/B) ratio in the microbiota of Indian subjects was investigated in all the subjects (n=1003). The obese individuals were also included in this analysis because there no strong effect of obesity status was observed on the microbiota community composition (see above). A continuum was detected irrespective of the BMI values where only a few subjects had exhibited high P/B ratio (Figure 6A). The subjects across geographies had a microbiota characterised by high P/B ratio (Figure 6B). Additionally, no significant correlation was observed between BMI and age with P/B ratio in the study cohort (Supplementary figure 3). The differences of P/B ratio between genders (male/female) was also not significant (Supplementary figure 4). P/B ratio showed significant correlation with the PCoA axis 1 which explained 30.6% of the variation in the microbial community in the study cohort (Supplementary figure 5).

**Figure 6:**
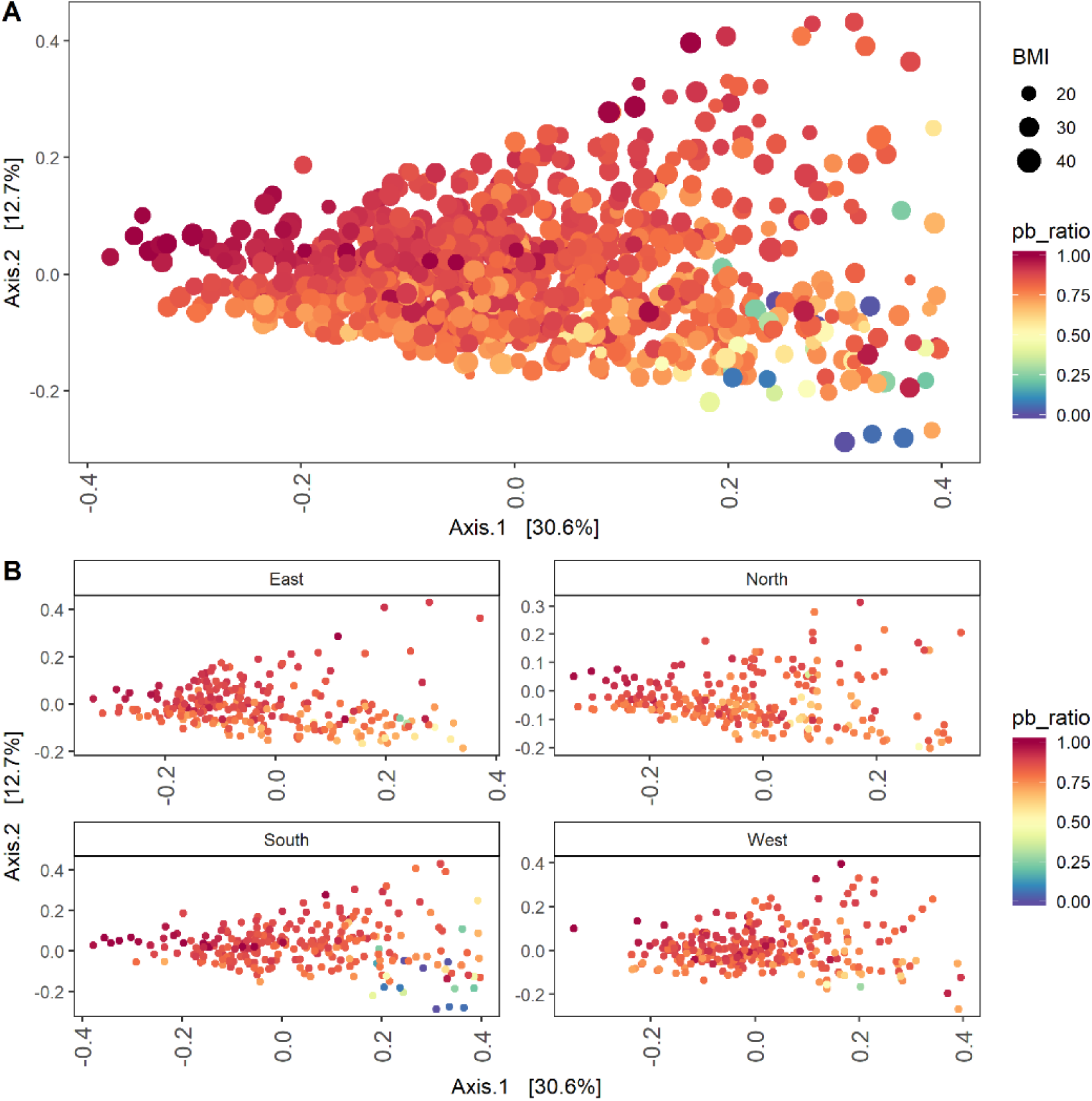
Principal coordinates analysis based on Bray-Curtis dissimilarity based on genus level relative abundances. **A]** PCoA depicting the gradient of *Prevotella/Bacteroides* ratio and distribution of body mass index (BMI) in 1003 Indian subjects. **B]** Same PCoA as in panel A, but coloured and facetted for depicting the distribution of *Prevotella/Bacteroides* ratio in the Indian gut microbiota in different geographical zones (East, n = 250; North, n = 243; South, n = 250; West, n = 260).

## Summary

The gut microbiome of Indian subjects differs in composition at phylum level across the four geographical zones. Overall variation in the gut microbial community structure in Indians is mostly driven by city of residence. Despite the large differences in the geographic location, there exists a core of 12 OTUs that are shared among 80% of the subjects. These core OTUs are classified as members of genera that are known for their ability to degrade complex polysaccharides (*Prevotella*, *Bacteroides*), produce butyrate and propionate (*Faecalibacterium*, *Megasphaera*) as well as ability to degrade mucin (*Ruminococcus gnavus*). Compared to the Westernized urban populations, the Indian population still harbours features of non-industrialized gut microbiota such as *Treponema*, which was present in 73% of the subjects at a low mean relative abundance of 0.004. Previously, *Treponema* was found to be characteristic of a traditional microbiome (Obregon-Tito *et al.*, 2015). Therefore, efforts need to made for cultivating and preserving human gut origin *Treponema* isolates from diverse populations that are undergoing rapid transition towards a Western lifestyle. The majority of variation in the microbial community structure was correlated with the ratio of *Prevotella* versus *Bacteroides*. However, due to lack of information on dietary habits, no concrete associations could be made to explain the high P/B ratio observed in the Indian population. Since both *Bacteroides* and *Prevotella* are capable of degrading complex polysaccharides, there is need to identify the trade-off between *Prevotella* or *Bacteroides* domination in the Westernized and urban Indian gut microbiota.

## Methods

### Data from LogMPIE

The data analysed in this study was obtained from figshare (Dubey *et al.*, 2018). Detailed information on sample collection and processing for DNA extraction, 16S rRNA gene amplification and sequencing are provided in the original publication (Dubey *et al.*, 2018). Here, some key points are described. The samples were collected by participants using sterile OMNIgene®•GUT stool collection kit. DNA extraction was done using the QiaAmp DNA Stool Mini Kit (Qiagen, Hilden, Germany). Two primer pairs one of V3 and one for V4 hypervariable region of the 16S rRNA gene was used for amplification (Milani *et al.*, 2013, Dubey *et al.*, 2018). The sequencing was done Ion S5 System (Thermo Fisher Scientific, Carlsbad, CA, USA). OTU tables were obtained by processing raw reads following the QIIME workflow on the Ion Reporter Server. OTU picking was done using the *pick_closed_reference_otus.py* command in QIIME (Caporaso *et al.*, 2010).

### Microbial community data handling, analysis and visualisation

The relative abundance microbial profiling data and metadata were obtained from (Dubey *et al.*, 2018)(https://doi.org/10.6084/m9.figshare.c.4147079.v1). The taxonomy was corrected to make it compatible with *read*_*phyloseq* function of microbiome R package (Lahti & Shetty, 2018). The resulting phyloseq object was analysed in R (v3.5.1) using the phyloseq (v1.24.1) and microbiome R package (v1.2.1) (McMurdie & Holmes, 2013, Lahti & Shetty, 2018). One subject, Subject-8032 was removed since initial ordinations revealed it to be highly divergent and thus the analysis was limited to 1003 subjects. Data visualisation was done using a combination of ggplot2 (v3.1) and ggpubr (v0.1.8) packages.

### Statistical analysis

The dissimilarity in gut microbiota composition between the subjects were investigated using the Bray-Curtis dissimilarity index calculated at OTU level and genus level relative abundance data using the phyloseq R package. The unconstrained principal coordinates analysis PCoA ordinations were visualised using the *plot*_*ordination* function. The contribution of each of the metadata categories, geographical location (city of residence), geographical zone, gender, and lifestyle pattern was investigated using PERMANOVA (999 permutations) (*adonis* function, vegan (v2.5-3) R package). Pair-wise comparisons were done using Wilcoxon test. Correlations between Prevotella versus Bacteroides ratios with age, BMI and PCoA axis 1 were based on Pearson’s correlations and done using the *stat_cor* function and visualised using *ggscater* function in ggpubr.

### Core microbiota analysis

The core microbiota analysis was done using the blanket approach (Salonen *et al.*, 2012). In this approach, the random sub-samples are drawn and the frequency of an OTU to be present in user defined samples (here, 810 samples) at a minimum relative abundance threshold (here, 0.0001) was calculated. Using 1000 bootstrap (boot, v1.3-20, R package) the median core size was estimated (Canty & Ripley, 2012). The effect of prevalence and abundance thresholds as well as the abundance and prevalence distribution of core OTUs were visualized using the microbiome R package (Lahti & Shetty, 2018).

### Prevotella/Bacteroides ratio analysis

The *Prevotella*/*Bacteroides* ratio were analysed using the approach described previously (Gorvitovskaia *et al.*, 2016). The OTU data was aggregated at genus level and used for further analysis.

## Supporting information

## Acknowledgement

The author would like to thank Ashok Kumar Dubey, Niyati Uppadhyaya, Pravin Nilawe, Neeraj Chauhan, Santosh Kumar, Urmila Anurag Gupta and Anirban Bhaduri, the authors of the “Landscape Of Gut Microbiome - Pan-India Exploration”, or LogMPIE study for making the data free and openly accessible. Without LogMPIE data this study would not have been possible.

## Funding

SAS is employed by the Laboratory of Microbiology Wageningen University and Research. This research was partly funded by the Netherlands Organisation for Scientific Research (NOW) Soehngen Institute of Anaerobic Microbiology (SIAM) grant and the NWO UNLOCK grant. The funders had no role in design and interpretation of this study.

## Conflict of interest

The author declares no conflict of interest.

## Data and code availability

The raw sequencing files are made available by the authors at European nucleotide archive (ENA) under the primary accession code, PRJEB25642, and secondary accession code, ERP07577 (Dubey *et al.*, 2018). The codes used for the analysis done in this manuscript will be made available at the following GitHub repository (https://github.com/microsud/Indian-gut-microbiota).

## Supplementary data

**Supplementary figure 1:**
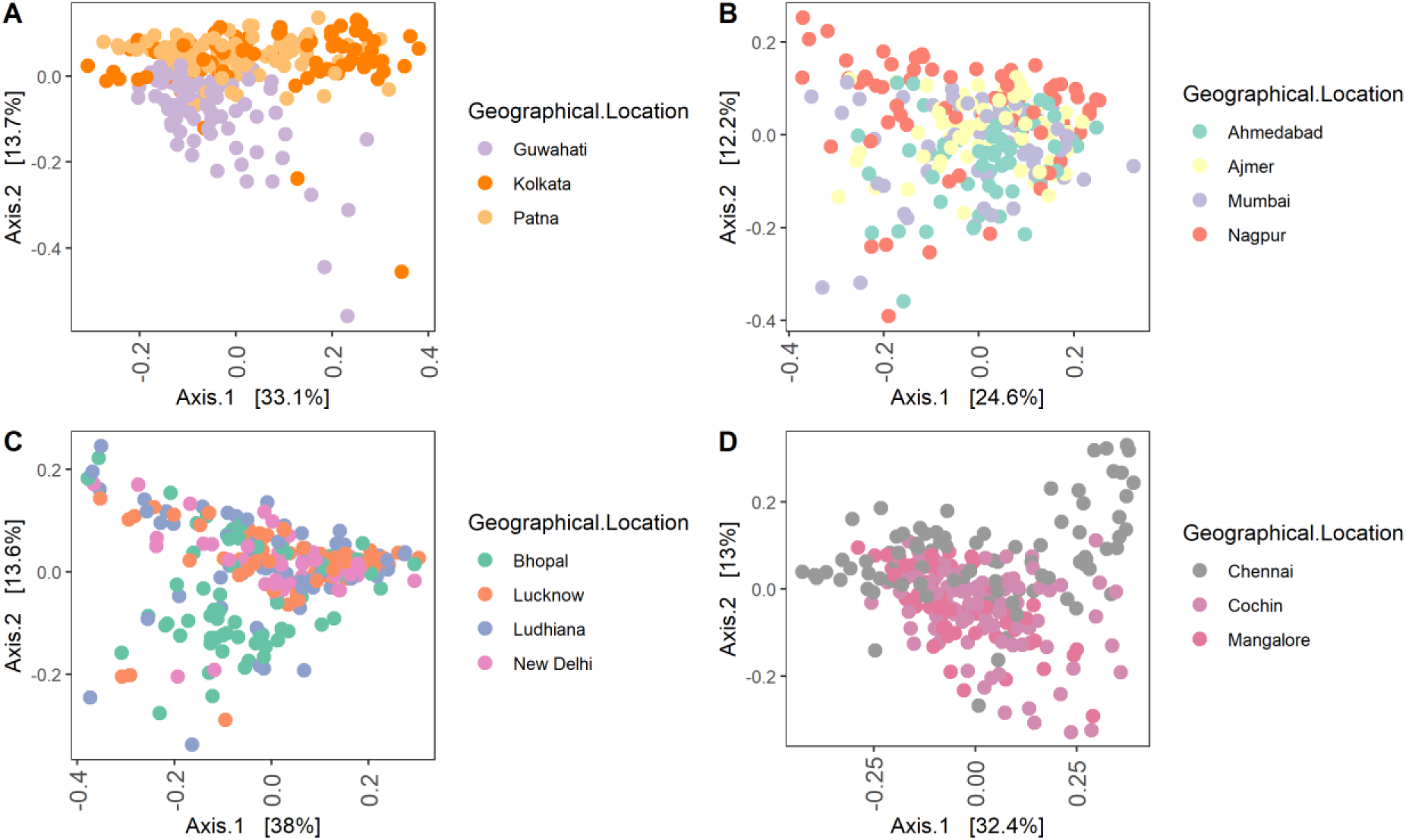
Principal coordinates analysis (PCoA) based on Bray-Curtis dissimilarity based on genus-level relative abundances. **A]** East; **B]** West; **C]** North **D]** South.

**Supplementary figure 2:**
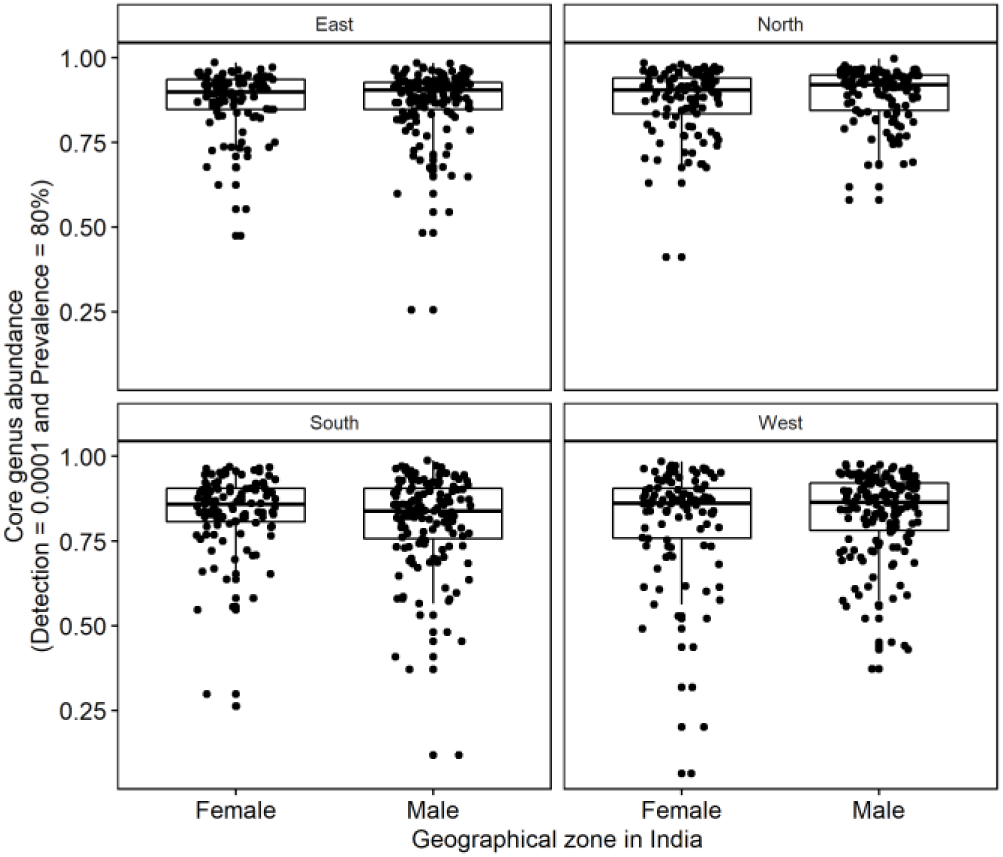
Contribution of 13 core genera towards the total abundance in the Indian female (n = 419) and male (n = 584) gut microbiota. The core microbiota was determined based on a minimum relative abundance of 0.0001 in present in minimum of 80% subjects (1000 bootstraps). The core genera were *Prevotella*, *Faecalibacterium*, *Bacteroides*, *Eubacterium*, *Roseburia*, *Ruminococcus*, *Lactobacillus*, *Megasphaera*, *Sutterella*, *Gemmiger*, *Blautia*, *Clostridium*, *Dorea*

**Supplementary figure 3:**
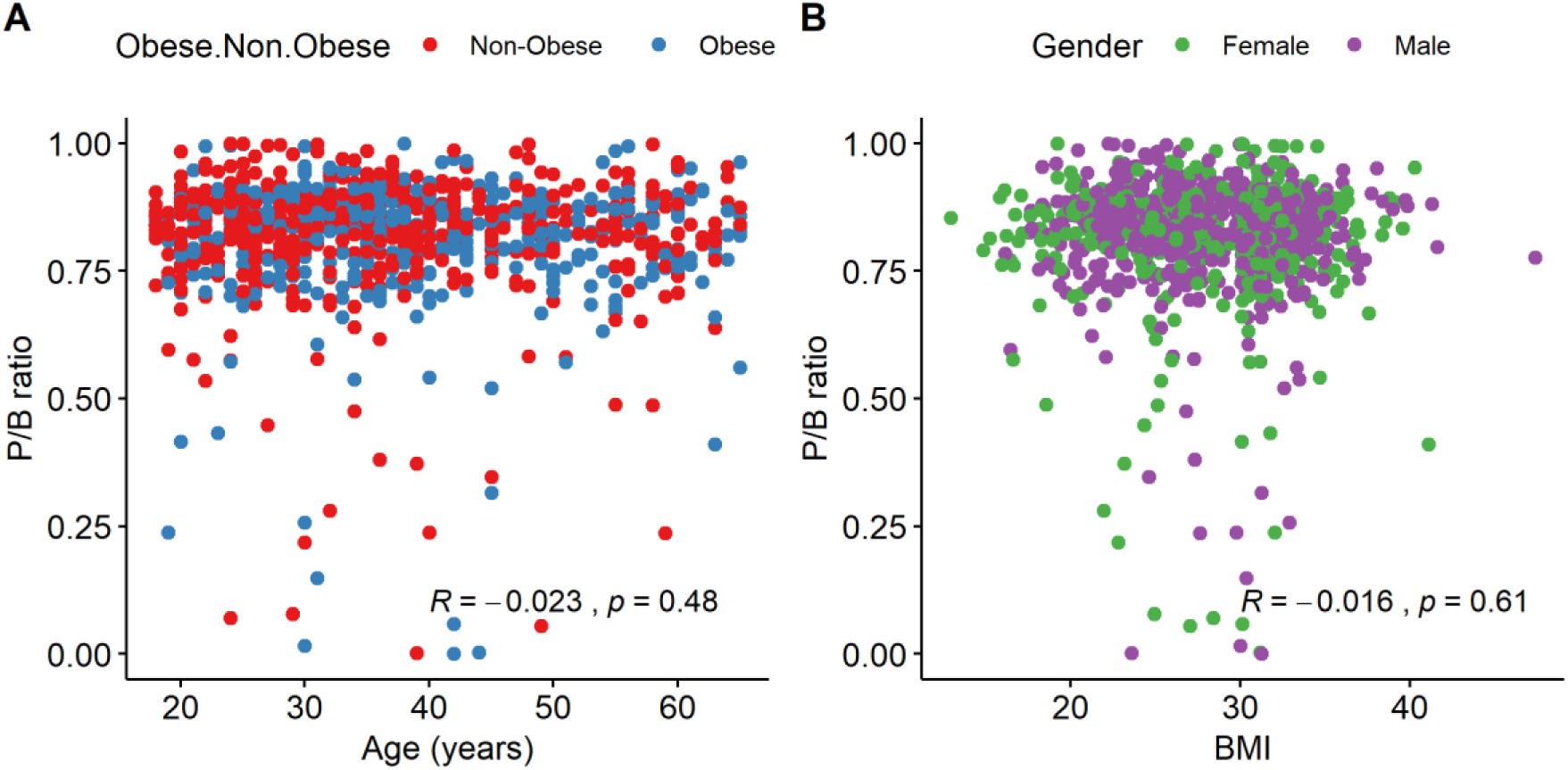
Pearson’s correlation analysis. **A]** Relationship between *Prevotella*/*Bacteroides* ratio and Age. **B]** Relationship between *Prevotella*/*Bacteroides* ratio and body mass index (BMI).

**Supplementary figure 4:**
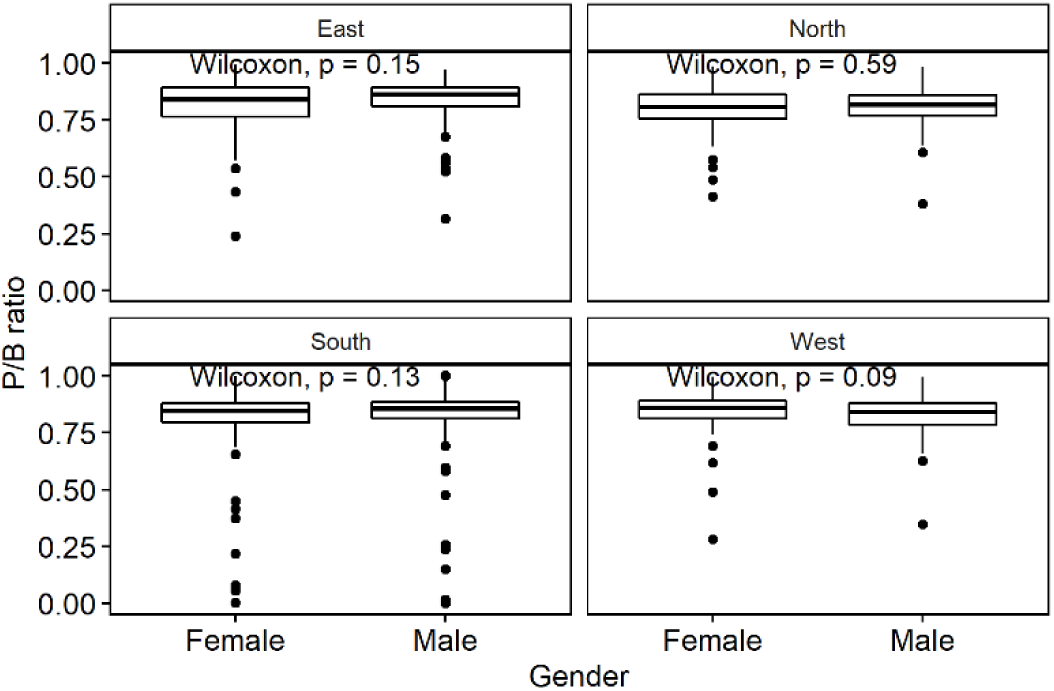
Comparison of P/B ratio between genders from different geographical locations. The p-values were calculated using Wilcoxon test.

**Supplementary figure 5:**
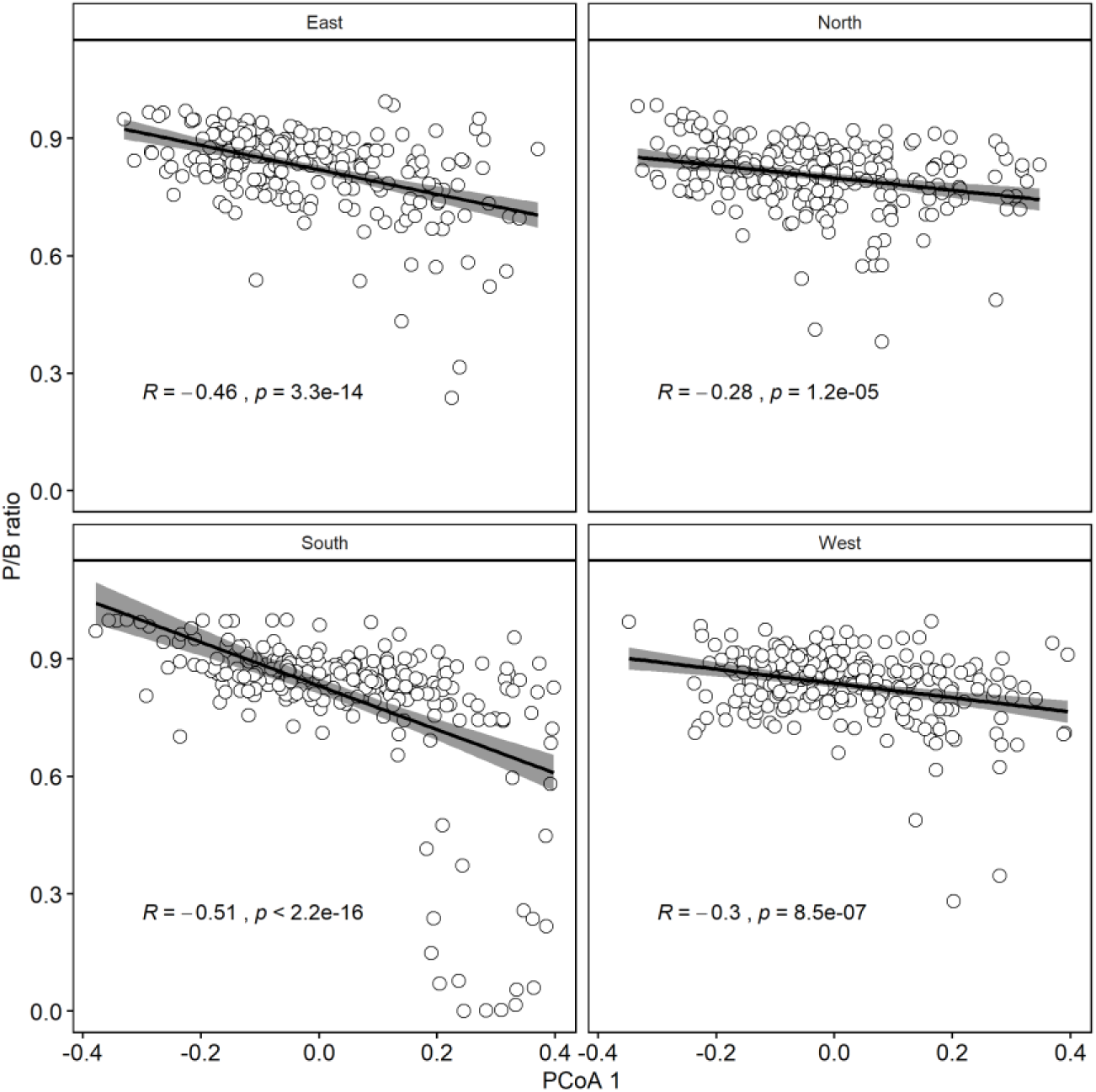
Pearson’s correlation between PCoA axis 1 and P/B ratio.

## References

Barcenilla A, Pryde SE, Martin JC, Duncan SH, Stewart CS, Henderson C & Flint HJ (2000) Phylogenetic relationships of butyrate-producing bacteria from the human gut. Applied and environmental microbiology 66: 1654–1661.

Baxter NT, Ruffin MT, Rogers MA & Schloss PD (2016) Microbiota-based model improves the sensitivity of fecal immunochemical test for detecting colonic lesions. Genome medicine 8: 37.

Bhute S, Pande P, Shetty SA, Shelar R, Mane S, Kumbhare SV, Gawali A, Makhani H, Navandar M & Dhotre D (2016) Molecular characterization and meta-analysis of gut microbial communities illustrate enrichment of Prevotella and Megasphaera in Indian subjects. Frontiers in microbiology 7: 660.

Canty A & Ripley B (2012) boot: Bootstrap R (S-Plus) functions. R package version 1.

Caporaso JG, Kuczynski J, Stombaugh J, Bittinger K, Bushman FD, Costello EK, Fierer N, Pena AG, Goodrich JK & Gordon JI (2010) QIIME allows analysis of high-throughput community sequencing data. Nature methods 7: 335.

Cotillard A, Kennedy SP, Kong LC, Prifti E, Pons N, Le Chatelier E, Almeida M, Quinquis B, Levenez F & Galleron N (2013) Dietary intervention impact on gut microbial gene richness. Nature 500: 585–588.

Dao MC, Everard A, Aron-Wisnewsky J, Sokolovska N, Prifti E, Verger EO, Kayser BD, Levenez F, Chilloux J & Hoyles L (2015) Akkermansia muciniphila and improved metabolic health during a dietary intervention in obesity: relationship with gut microbiome richness and ecology. Gut gutjnl-2014-308778.

Das B, Ghosh TS, Kedia S, Rampal R, Saxena S, Bag S, Mitra R, Dayal M, Mehta O & Surendranath A (2018) Analysis of the gut microbiome of rural and urban healthy indians living in sea level and high altitude areas. Scientific reports 8: 10104.

David LA, Maurice CF, Carmody RN, Gootenberg DB, Button JE, Wolfe BE, Ling AV, Devlin AS, Varma Y & Fischbach MA (2014) Diet rapidly and reproducibly alters the human gut microbiome. Nature 505: 559–563.

De Filippo C, Cavalieri D, Di Paola M, Ramazzotti M, Poullet JB, Massart S, Collini S, Pieraccini G & Lionetti P (2010) Impact of diet in shaping gut microbiota revealed by a comparative study in children from Europe and rural Africa. Proceedings of the National Academy of Sciences 107: 14691–14696.

Dehingia M, Talukdar NC, Talukdar R, Reddy N, Mande SS, Deka M & Khan MR (2015) Gut bacterial diversity of the tribes of India and comparison with the worldwide data. Scientific reports 5: 18563.

Delgado-Baquerizo M, Oliverio AM, Brewer TE, Benavent-González A, Eldridge DJ, Bardgett RD, Maestre FT, Singh BK & Fierer N (2018) A global atlas of the dominant bacteria found in soil. Science 359: 320–325.

Desai MS, Seekatz AM, Koropatkin NM, Kamada N, Hickey CA, Wolter M, Pudlo NA, Kitamoto S, Terrapon N & Muller A (2016) A dietary fiber-deprived gut microbiota degrades the colonic mucus barrier and enhances pathogen susceptibility. Cell 167: 1339–1353. e1321.

Dubey AK, Uppadhyaya N, Nilawe P, Chauhan N, Kumar S, Gupta UA & Bhaduri A (2018) LogMPIE, pan-India profiling of the human gut microbiome using 16S rRNA sequencing. Scientific Data 5: 180232.

Everard A, Belzer C, Geurts L, Ouwerkerk JP, Druart C, Bindels LB, Guiot Y, Derrien M, Muccioli GG & Delzenne NM (2013) Cross-talk between Akkermansia muciniphila and intestinal epithelium controls diet-induced obesity. Proceedings of the National Academy of Sciences 110: 9066–9071.

Falony G, Joossens M, Vieira-Silva S, et al. (2016) Population-level analysis of gut microbiome variation. Science 352: 560–564.

Flint HJ, Scott KP, Louis P & Duncan SH (2012) The role of the gut microbiota in nutrition and health. Nature Reviews Gastroenterology and Hepatology 9: 577.

Flint HJ, Scott KP, Duncan SH, Louis P & Forano E (2012) Microbial degradation of complex carbohydrates in the gut. Gut microbes 3: 289–306.

Ghosh TS, Gupta SS, Nair GB & Mande SS (2013) In silico analysis of antibiotic resistance genes in the gut microflora of individuals from diverse geographies and age-groups. PLoS One 8: e83823.

Gorvitovskaia A, Holmes SP & Huse SM (2016) Interpreting Prevotella and Bacteroides as biomarkers of diet and lifestyle. Microbiome 4: 1.

Hayashi H, Shibata K, Sakamoto M, Tomita S & Benno Y (2007) Prevotella copri sp. nov. and Prevotella stercorea sp. nov., isolated from human faeces. International journal of systematic and evolutionary microbiology 57: 941–946.

Heinken A, Khan MT, Paglia G, Rodionov DA, Harmsen HJM & Thiele I (2014) Functional metabolic map of *Faecalibacterium prausnitzii*, a beneficial human gut microbe. Journal of Bacteriology 196: 3289–3302.

Hosseini E, Grootaert C, Verstraete W & Van de Wiele T (2011) Propionate as a health-promoting microbial metabolite in the human gut. Nutrition Reviews 69: 245–258.

Human Microbiome Project C (2012) Structure, function and diversity of the healthy human microbiome. Nature 486: 207–214.

Huse SM, Ye Y, Zhou Y & Fodor AA (2012) A core human microbiome as viewed through 16S rRNA sequence clusters. PloS one 7: e34242.

Huttenhower C, Gevers D, Knight R, Abubucker S, Badger JH, Chinwalla AT, Creasy HH, Earl AM, FitzGerald MG & Fulton RS (2012) Structure, function and diversity of the healthy human microbiome. Nature 486: 207.

Jalanka-Tuovinen J, Salonen A, Nikkilä J, Immonen O, Kekkonen R, Lahti L, Palva A & de Vos WM (2011) Intestinal microbiota in healthy adults: temporal analysis reveals individual and common core and relation to intestinal symptoms. PloS one 6: e23035.

Jia X, Dini-Andreote F & Salles JF (2018) Community Assembly Processes of the Microbial Rare Biosphere. Trends in microbiology.

Lahti L & Shetty SA (2018) Tools for microbiome analysis in R.

Lahti L, Salojärvi J, Salonen A, Scheffer M & de Vos WM (2014) Tipping elements in the human intestinal ecosystem. Nat Commun 5.

Li J, Jia H, Cai X, Zhong H, Feng Q, Sunagawa S, Arumugam M, Kultima JR, Prifti E & Nielsen T (2014) An integrated catalog of reference genes in the human gut microbiome. Nat Biotech 32: 834–841.

Lin HV, Frassetto A, Kowalik Jr EJ, Nawrocki AR, Lu MM, Kosinski JR, Hubert JA, Szeto D, Yao X & Forrest G (2012) Butyrate and propionate protect against diet-induced obesity and regulate gut hormones via free fatty acid receptor 3-independent mechanisms. PloS one 7: e35240.

Louis P & Flint HJ (2017) Formation of propionate and butyrate by the human colonic microbiota. Environmental microbiology 19: 29–41.

Lynch MD & Neufeld JD (2015) Ecology and exploration of the rare biosphere. Nature Reviews Microbiology 13: 217.

Martínez I, Stegen JC, Maldonado-Gómez MX, Eren AM, Siba PM, Greenhill AR & Walter J (2015) The gut microbiota of rural papua new guineans: composition, diversity patterns, and ecological processes. Cell reports 11: 527–538.

McMurdie PJ & Holmes S (2013) phyloseq: an R package for reproducible interactive analysis and graphics of microbiome census data. PloS one 8: e61217.

Milani C, Hevia A, Foroni E, Duranti S, Turroni F, Lugli GA, Sanchez B, Martin R, Gueimonde M & Van Sinderen D (2013) Assessing the fecal microbiota: an optimized ion torrent 16S rRNA gene-based analysis protocol. PloS one 8: e68739.

O’Keefe SJD, Li JV, Lahti L, et al.(2015) Fat, fibre and cancer risk in African Americans and rural Africans. Nat Commun 6: 6342.

Obregon-Tito AJ, Tito RY, Metcalf J, Sankaranarayanan K, Clemente JC, Ursell LK, Xu ZZ, Van Treuren W, Knight R & Gaffney PM (2015) Subsistence strategies in traditional societies distinguish gut microbiomes. Nat Commun 6: 6505.

Plovier H, Everard A, Druart C, Depommier C, Van Hul M, Geurts L, Chilloux J, Ottman N, Duparc T & Lichtenstein L (2017) A purified membrane protein from Akkermansia muciniphila or the pasteurized bacterium improves metabolism in obese and diabetic mice. Nature medicine 23: 107.

Qin J, Li R, Raes J, Arumugam M, Burgdorf KS, Manichanh C, Nielsen T, Pons N, Levenez F & Yamada T (2010) A human gut microbial gene catalogue established by metagenomic sequencing. nature 464: 59–65.

Qin J, Li Y, Cai Z, et al.(2012) A metagenome-wide association study of gut microbiota in type 2 diabetes. Nature 490: 55–60.

Reichardt N, Duncan SH, Young P, Belenguer A, Leitch CM, Scott KP, Flint HJ & Louis P (2014) Phylogenetic distribution of three pathways for propionate production within the human gut microbiota. The ISME journal 8: 1323.

Rothschild D, Weissbrod O, Barkan E, Kurilshikov A, Korem T, Zeevi D, Costea PI, Godneva A, Kalka IN & Bar N (2018) Environment dominates over host genetics in shaping human gut microbiota. Nature 555: 210.

Salonen A, Salojärvi J, Lahti L & De Vos W (2012) The adult intestinal core microbiota is determined by analysis depth and health status. Clinical Microbiology and Infection 18: 16–20.

Schubert AM, Rogers MA, Ring C, Mogle J, Petrosino JP, Young VB, Aronoff DM & Schloss PD (2014) Microbiome data distinguish patients with Clostridium difficile infection and non-C. difficile-associated diarrhea from healthy controls. MBio 5: e01021–01014.

Shade A, Jones SE, Caporaso JG, Handelsman J, Knight R, Fierer N & Gilbert JA (2014) Conditionally rare taxa disproportionately contribute to temporal changes in microbial diversity. MBio 5: e01371–01314.

Shetty SA, Marathe NP & Shouche YS (2013) Opportunities and challenges for gut microbiome studies in the Indian population. Microbiome 1: 24.

Shetty SA, Hugenholtz F, Lahti L, Smidt H & de Vos WM (2017) Intestinal microbiome landscaping: insight in community assemblage and implications for microbial modulation strategies. FEMS microbiology reviews 41: 182–199.

Sonnenburg ED, Zheng H, Joglekar P, Higginbottom SK, Firbank SJ, Bolam DN & Sonnenburg JL (2010) Specificity of polysaccharide use in intestinal bacteroides species determines diet-induced microbiota alterations. Cell 141: 1241–1252.

Tandon D, Haque MM, Saravanan R, Shaikh S, Sriram P, Dubey AK & Mande SS (2018) A snapshot of gut microbiota of an adult urban population from Western region of India. PloS one 13: e0195643.

Turnbaugh PJ, Hamady M, Yatsunenko T, Cantarel BL, Duncan A, Ley RE, Sogin ML, Jones WJ, Roe BA & Affourtit JP (2009) A core gut microbiome in obese and lean twins. nature 457: 480–484.

Yajnik CS & Yudkin JS (2004) The YY paradox. The Lancet 363: 163.

Yano JM, Yu K, Donaldson GP, Shastri GG, Ann P, Ma L, Nagler CR, Ismagilov RF, Mazmanian SK & Hsiao EY (2015) Indigenous bacteria from the gut microbiota regulate host serotonin biosynthesis. Cell 161: 264–276.

Yatsunenko T, Rey FE, Manary MJ, Trehan I, Dominguez-Bello MG, Contreras M, Magris M, Hidalgo G, Baldassano RN & Anokhin AP (2012) Human gut microbiome viewed across age and geography. Nature 486: 222–227.

Zeller G, Tap J, Voigt AY, Sunagawa S, Kultima JR, Costea PI, Amiot A, Böhm J, Brunetti F & Habermann N (2014) Potential of fecal microbiota for early-stage detection of colorectal cancer. Molecular systems biology 10: 766.

Zhang J, Guo Z, Lim AAQ, Zheng Y, Koh EY, Ho D, Qiao J, Huo D, Hou Q & Huang W (2014) Mongolians core gut microbiota and its correlation with seasonal dietary changes. Scientific reports 4.

Zhang J, Guo Z, Lim AAQ, Zheng Y, Koh EY, Ho D, Qiao J, Huo D, Hou Q & Huang W (2014) Mongolians core gut microbiota and its correlation with seasonal dietary changes. Scientific reports 4: 5001.

